# Cytonuclear Conflict and Reticulate Evolution in the Morelloid Clade (*Solanum*, Solanaceae): Insights from Genome Skimming and Network Phylogenomics

**DOI:** 10.64898/2026.02.26.708157

**Authors:** Sundre Winslow, Sandra Knapp, Tiina Särkinen, Péter Poczai

**Affiliations:** Botany and Mycology Unit, Finnish Museum of Natural History & Department of Organismal and Evolutionary Biology, Faculty of Biological and Environmental Sciences PO Box 7 University of Helsinki, FI-00014 Unioninkatu 44, Helsinki, Finland; Natural History Museum, Cromwell Road, London SW7 5BD United Kingdom; Royal Botanic Garden Edinburgh, 20A Inverleith Row, Edinburgh EH3 5LR, United Kingdom

**Author notes:** Corresponding author: Sundre Winslow; Péter Poczai.

**Keywords:** Phylogenomics, Solanum, Solanaceae, Reticulate evolution, Cytonuclear discordance, Herbarium genomics, Genome skimming

## Abstract

The Morelloid clade (black nightshades) is one of the most strongly supported clades within the megadiverse *Solanum* genus. It comprises 76 globally distributed, non-spiny herbaceous and suffrutescent species. While often erroneously considered poisonous weeds, several species are economically important as orphan crops. The clade is closely related to tomato and potato but, due to a lack of focused breeding efforts, remains a reservoir of genetic diversity for crop improvement. Despite this potential, we lack fundamental knowledge on the evolution of the Morelloid clade. The group includes polyploid species with unknown parental origins—likely reflecting reticulate processes such as hybridization, introgression, and associated backcrossing events. Prior analyses have been unable to disentangle these processes, leaving the mechanisms underlying reticulate evolution in the Morelloid clade poorly understood. Here, we use genome skimming to produce a well-supported maximum likelihood plastid phylogeny from complete circularized plastomes and a coalescent-based species tree from combined Angiosperms353 and conserved ortholog set nuclear markers. Our dataset, composed of previously published data and deep genome skimming from herbarium samples, spans 26 Morelloid species. To investigate patterns of non-treelike evolution, we used a nuclear phylogenetic network, multispecies coalescent simulations, a fused rooted nuclear chloroplast tree, and quantification of nuclear gene tree concordance. We show that incongruence between nuclear and plastid trees is pervasive and cannot be explained by incomplete lineage sorting alone. Instead, our results demonstrate that events consistent with repeated chloroplast capture have shaped the reticulate evolutionary history of the clade, especially among African polyploid and Pan-American diploid lineages.

## 1. Introduction

The genus *Solanum* L. is the largest genus within the Solanaceae, comprising around 1,400 species, and placing it among the 86 taxonomically stable “big plant genera” (Moonlight et al., 2024). Early efforts to classify *Solanum* relied predominantly on morphological traits such as the presence or absence of prickles and anther morphology (Dillenius, 1732; Linnaeus, 1753; Dunal, 1816). These systems were synthesized into D’Arcy’s (1972) widely used scheme, which recognized seven *Solanum* subgenera. However, the first large-scale phylogenetic analysis integrating molecular and morphological data later identified twelve major *Solanum* clades (Bohs, 2005). One of the most strongly supported was the Morelloid clade, a group not previously recognized based on morphology alone, that included the type of the genus (*S. nigrum* L.). The name for the group was derived from “Maurella,” which Dunal (1816) used to describe an unranked group of 15 species related to *S. nigrum*.

The Morelloid clade, as circumscribed now based on more detailed phylogenetic and taxonomic studies, comprises approximately 78 globally distributed species, with their center of diversity in South America (Gagnon et al., 2022; Knapp et al., 2019, 2023; Särkinen et al., 2018). The clade is characterized primarily by non-spiny herbaceous plants, though occasionally woody shrubs, with mostly internodal inflorescences and brightly colored, juicy berries (Knapp et al., 2019, 2023; Särkinen et al., 2018; Fig. 1). Their ploidy ranges from diploid to hexaploid. Many species show highly variable leaf morphology even within a single plant. As a result, there are 594 associated names assigned to the Morelloids (Knapp et al., 2019, 2023; Särkinen et al., 2018).

**Fig. 1.**
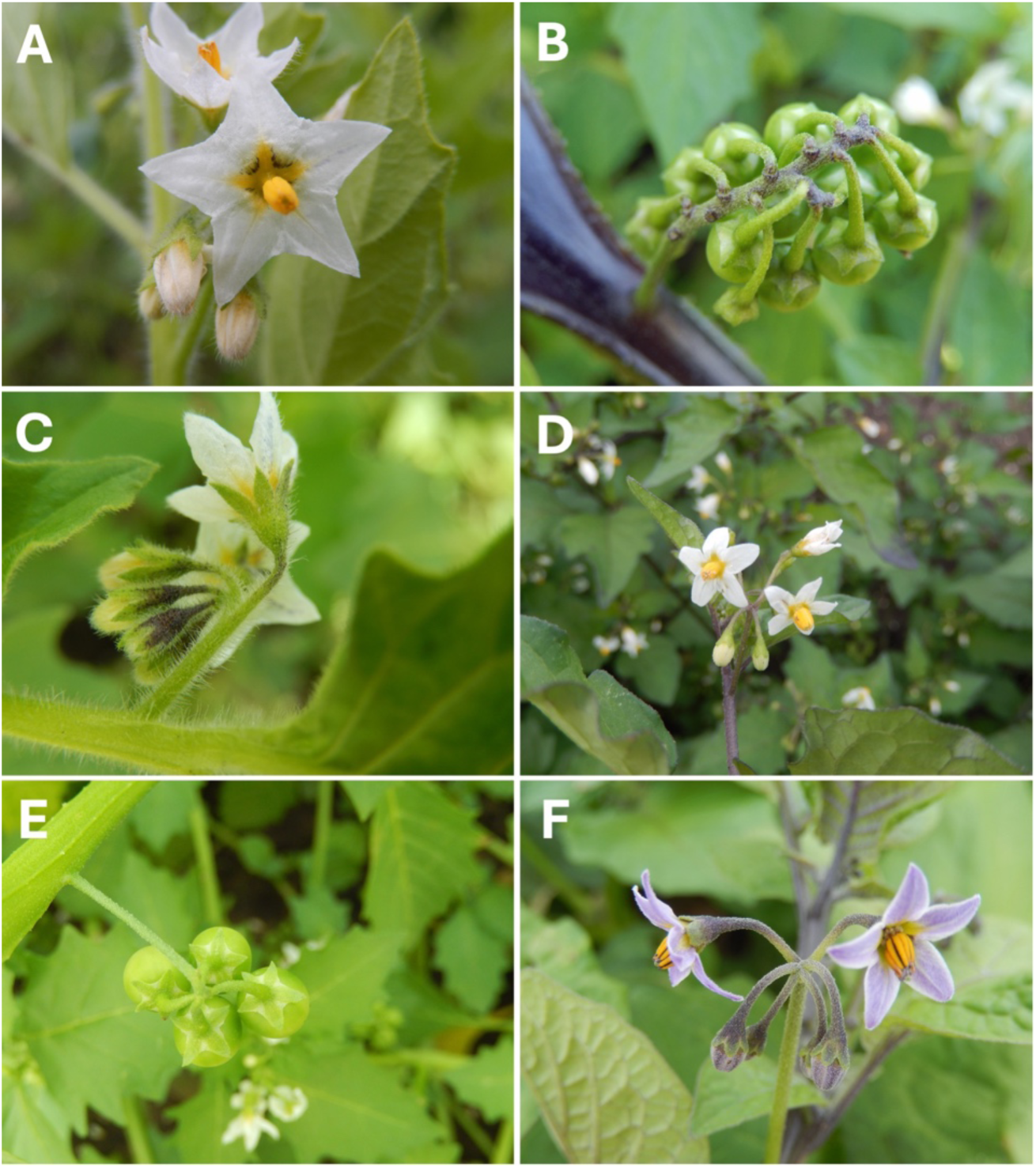
Morphology of the Morelloid clade of *Solanum*. A, Flowers of *S. physalifolium* Rusby; B, Berries of *S. umalilaense* Manoko; C, Buds and flowers of *S. sarrachoides* Sendtn.; D, Flowers of *S. nigrum* L.; E, Berries of *S. retroflexum* Dunal; F, Flowers and buds of *S. scabrum* Mill..

Several Morelloid species are of agricultural importance as orphan crops (Sogbohossou et al., 2018). *Solanum scabrum*, *S. villosum*, and *S. retroflexum* are commonly cultivated leafy vegetables in Africa and contribute to local food security and income. Beyond their direct use, Morelloids are a potential genetic reservoir for closely related crops such as potato and tomato. For example, *S. nigrum* and *S. americanum* have resistance genes against *Phytophthora infestans* (Mont.) de Bary, which have been used to produce stable transgenic lines in closely related species (Lebecka, 2009; Lin et al., 2022; Witek et al., 2016).

Polyploidy in *Solanum* is phylogenetically clustered, especially allopolyploidy, and is only observed in three *Solanum* clades: the Petota minor, Peregrina, and Morelloid clades (de Queiroz and Cantino, 2020). Discordant gene trees between plastid and nuclear datasets suggest Morelloid polyploids have an allopolyploid origin (Särkinen et al., 2015). The parental origins of these polyploids remain unresolved, with preliminary analysis implicating *S. chenopodioides*, *S. nitidibaccatum*, *S. americanum,* and *S. villosum* as putative parental species (Poczai & Hyvönen, 2011). Results largely confirm the non-hierarchical clustering analysis by Edmonds (1978).

Together, these patterns suggest that evolutionary histories within the Morelloid clade may not always conform to strictly bifurcating models.

The *S. tarderemotum* complex has posed taxonomic challenges, with disagreement over whether *S. florulentum* represents a distinct species or is a morphological variant of *S. tarderemotum*. Olet (2004) described three morphological forms of *S. tarderemotum* and recognized *S. florulentum* as a separate species, whereas Särkinen et al. (2018) argue that they are not morphologically distinct. Manoko (2024) observed five clusters within an amplified fragment length polymorphisms (AFLP)-based phenetic tree. By comparing morphology with genetic data, they recovered two distinct clusters corresponding to *S. florulentum* and *S. tarderemotum*, respectively, with three intermediary hybrid groups. In four cases, individuals grown from seeds from the same mother plant were found in different clusters, which Manoko (2024) attributed to potential F₂ hybrids. This could be evidence of interspecific hybridization within the *S. tarderemotum* complex, though these methods only provide preliminary results.

The Morelloids exhibit extensive morphological variation and often have unclear species boundaries. Lin et al. (2022) generated a phylogeny of 85 *S. americanum* accessions, identifying four distinct clades. Six additional accessions fell outside these clades and were considered outliers. Four of these outlier accessions (SP2303, SP2310, SP3393, and SP3052) exhibited high heterozygosity, suggesting polyploidy or introgression. The other two accessions (SP3052 and SP3376) appear to represent a different diploid *Solanum* species than *S. americanum*. The unexpected placement of these accessions underscores the limitations of morphology-based identification in the Morelloid clade and highlights unresolved relationships.

Increasingly, studies of plant evolution demonstrate that using methods that assume strictly bifurcating phylogenetic trees often fail to capture the complexity of lineages that have undergone hybridization, introgression, and polyploidy (Hodel et al., 2022; Nie et al., 2023; Persson et al., 2020; Talavera et al., 2023; Zecca et al., 2020). Cytonuclear discordance is a key indicator of such non-dichotomous histories, frequently reflecting chloroplast capture or asymmetric gene flow rather than incomplete lineage sorting alone (Rieseberg & Soltis, 1991; Stegemann et al., 2012; Tsitrone et al., 2003). In the Morelloid clade, discordance between plastid and nuclear datasets, together with extensive polyploidy and putative hybrid lineages, suggests that reticulate evolution may be pervasive. However, most previous studies have relied on limited loci or single-marker datasets, thereby limiting their ability to disentangle introgression from other processes, such as incomplete lineage sorting.

Here, we use genome skimming to extract complete plastomes and nuclear markers from 25 Morelloid species. This study aims to reconstruct both nuclear and plastid Morelloid phylogenies, to investigate cytonuclear discordance and reticulate evolution within the clade. Our data finds that the Morelloids have no clear dichotomous history. Instead, the clade is represented by a reticulate network, where historical chloroplast introgression may best explain the observed cytonuclear incongruence.

## 2. Materials and Methods

### 2.1 Taxon sampling

We sampled 25 of the ca. 76 species within the Morelloid clade, representing 12 South American, 6 African, and 2 North American species, as well as Pan-American, circumtropical, and Eurasian species (Table 1). To test interspecific relationships, we sampled seven *S. tarderemotum* accessions, corresponding to those used by Manoko (2024), with one from each of the five clusters Manoko (2024) identified (Table S2). We included two *S. nigrum* samples from China, and both cultivated and wild forms of *S. scabrum*. The sampling included the recently described *S. caatingae* (S34; Knapp & Särkinen, 2018), found in the caatinga biome of north-eastern Brazil. This species is morphologically similar to *S. americanum* but differs in its glandular pubescence of translucent trichomes and long anthers. It was described based on herbarium sheets but has not yet been included in any genomic studies. We also included a weedy black nightshade, which was observed in Louisiana spreading rapidly through sugarcane (Orgeron et al., 2018). This species matched a previously described morphologically distinct sample, but based on *ndh*F-*rpl3*2 data, was shown to share near identity with *S. nigrescens*. Here, it is referred to as *S.* sp. Schilling (S23).

**Table 1.**
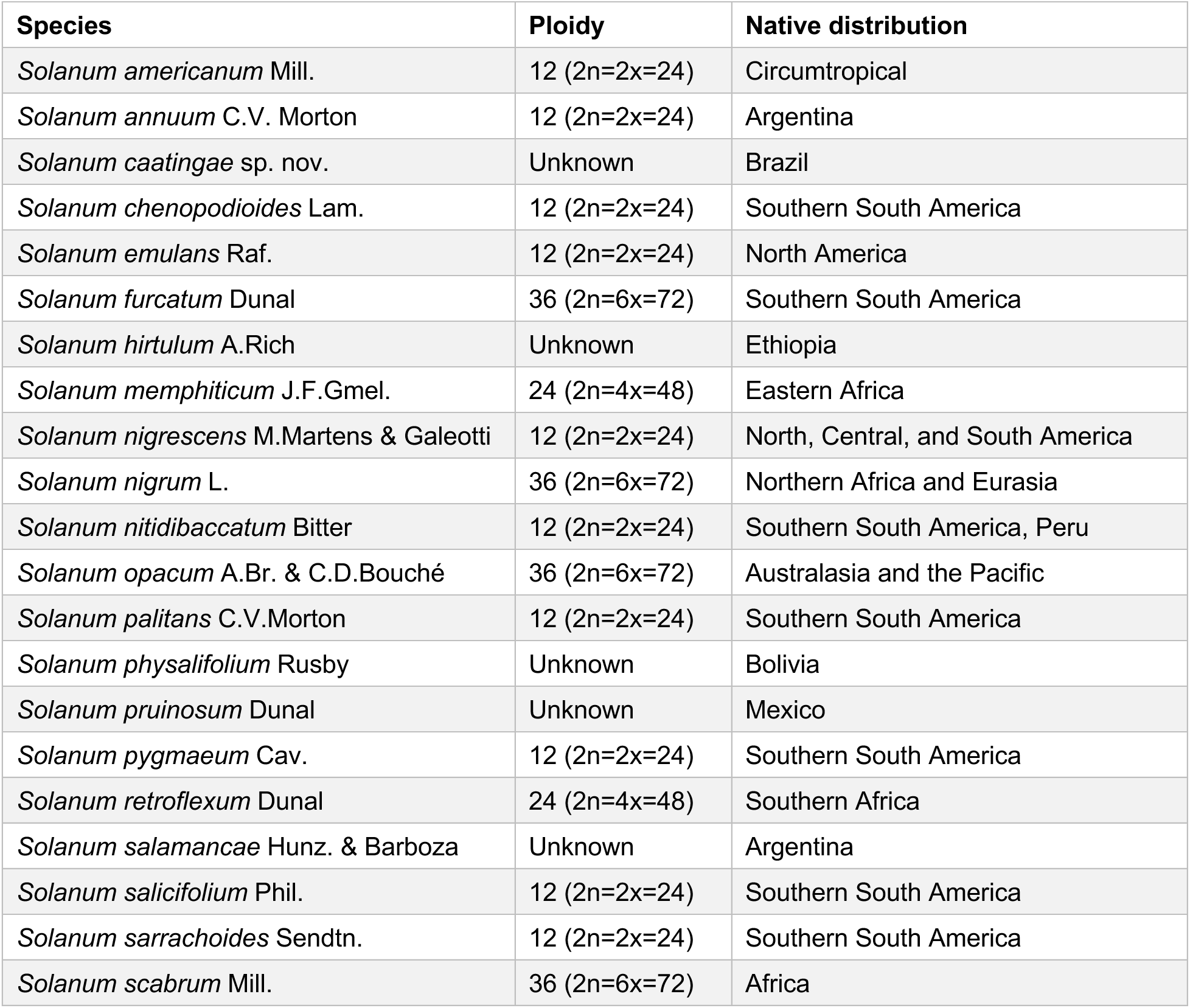
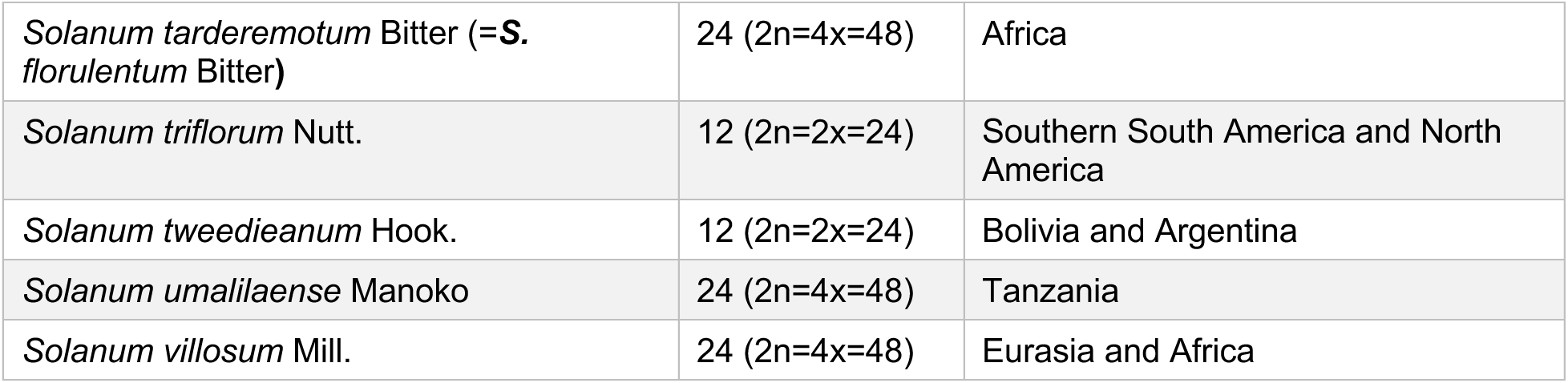
Species of the Morelloid clade of *Solanum* used in this study, including ploidy and distribution data.

To expand our sampling of *S. americanum*, we incorporated sequencing data generated by Lin et al. (2022). Genome-skimming data for 53 accessions were downloaded from the Sequence Read Archive (SRA) using SRAtools and converted to paired-end reads with fastq-dump. While all were classified as *S. americanum* in the Lin et al. (2022) publication, we have since studied the vouchers morphologically and identified many of the accessions considered “*S. americanum*” by the original authors as distinct species (Table S1). Revised species names reflecting these updated classifications were used throughout subsequent analyses. In addition, we used 18 assembled plastid genome sequences and the corresponding raw sequencing data for Morelloid taxa from Gagnon et al. (2022; Table S3). In total, this study examined 99 accessions, comprising 28 newly extracted sequences and 71 from published datasets.

### 2.2 Herbarium DNA extraction and deep genomes-skimming

We sampled 2–8 mm² leaf segments from herbarium specimens housed in the LUOMUS Botanical Collections (HEL), the Herbarium of the Royal Botanic Garden Edinburgh (E), and the Solanaceae Genbank Collection Nijmegen, as well as one specimen collected by Schilling in 2017 in Louisiana, USA (Table S2). DNA extractions were performed in the dedicated historical DNA (hDNA) laboratory at the Finnish Museum of Natural History to minimize the contamination of highly fragmented samples.

Extractions were carried out using either the DNeasy Plant Mini Kit (Qiagen, Venlo, The Netherlands) or the E.Z.N.A. Plant DNA DS Kit (Omega Bio-Tek, Norcross, GA, USA). DNA concentration and quality were assessed using a Qubit 4 fluorometer (Thermo Fisher Scientific, Waltham, MA, USA). DNA Integrity was evaluated using the TapeStation 4150 (Agilent Technologies, Santa Clara, CA, USA), which provides a DNA Integrity Number (DIN) to indicate degradation levels.

TruSeq DNA Nano libraries (Illumina, San Diego, CA, USA) were constructed from the herbarium DNA following the manufacturer’s protocol. Libraries were sequenced on a NovaSeq X Plus System with 2 × 150 bp paired-end reads. For each sample, we generated approximately 10 Gb of genome-skimming data.

### 2.3 Short-read data processing

We developed a contaminant database using genomes downloaded from NCBI (Sayers et al., 2019) and indexed with Bowtie2 (Langmead & Salzberg, 2012) (Bieker et al., 2020; Glassing et al., 2016). We used this database with FastQ Screen (Wingett & Andrews, 2018) to remove reads matching common herbarium and lab contaminants. Adapter sequences and low-quality bases were further removed using Trimmomatic v0.36 (Bolger et al., 2014), while six samples were additionally processed with FastP (Chen et al., 2018). The quality of the trimmed reads was assessed using FastQC (Andrews, 2010).

### 2.4 Chloroplast genome assembly

Plastome assemblies were generated using a combination of de novo and reference-guided methods across multiple software tools (Table S1 and S2). We used GetOrganelle v1.7.5 (Jin et al., 2020) with the embplant_pt seed database (-F), 15 extension iterations (-R), and default SPAdes k-mer settings (-k 21,45,65,85,105) to produce circular plastome assemblies. This approach successfully assembled twelve genomes.

For four genomes, circular assemblies were not obtained using the default embplant_pt seed. In these cases, we used the tomato plastome sequence (accession NC_007898.3) as a seed (-s) to improve assembly success. Nine more genomes were assembled using a seed-extended de novo approach via GeneMiner2 (Xie et al., 2024), which incorporated a custom reference database of published *Solanum* plastid genomes and internally utilized NOVOPlasty (Dierckxsens et al., 2017) with default settings.

Neither GetOrganelle nor NOVOPlasty produced a circular genome for *S. furcatum*, although read mapping to the tomato reference indicated good coverage and contiguity. To complete this assembly, we merged contigs generated by SPAdes in GetOrganelle using Geneious Prime 2024.0.5 (Biomatters Ltd, Auckland, New Zealand). assembler, then extended them further through iterative read mapping with Geneious’s built-in mapping tool. We manually inspected the resulting contigs, trimming low-confidence bases and replacing them with Ns as needed.

Final contigs were ordered using Mauve v2.3.1 (Darling et al., 2004; Rissman et al., 2009) against the tomato reference, followed by another round of iterative read mapping in Geneious to identify and correct remaining errors. Genome annotations were performed using GeSeq (Tillich et al., 2017), and circular maps were visualized with Chloroplot (Zheng et al., 2020). Inverted repeat junctions were further examined and plotted with IRplus (Menendez et al., 2023).

### 2.5 Chloroplast phylogeny

To generate a plastid-based phylogeny, genomes were aligned using MAFFT v7.49 (Katoh & Standley, 2013) within Geneious. One inverted repeat (IR) region was excluded to reduce overrepresentation in subsequent analyses. The alignment was visually inspected and manually adjusted at the 5′ end by removing gapped positions, then exported to FASTA format, which was used as the input for Bad2Matrix (Salinas et al., 2024). We implemented a motif-based adjustment protocol as proposed by Roestel et al. (2024).

Indel coding was applied following Simmons & Ochoterena (2000), since incorporating indels has been shown to enhance resolution and increase support for problematic nodes in phylogenetic inference (Donath & Stadler, 2018; Suvorov et al., 2022). We removed all remaining gapped positions and partitioned the dataset in Geneious by annotating coding and intergenic spacer (IGS) regions in the alignment. All partitions were extracted, and 34 regions (3,359 bp) containing only monomorphic sites were discarded. The remaining partitions were realigned individually with MAFFT using a custom batch workflow.

The filtered alignments were concatenated in Bad2Matrix using the -i flag to disable indel coding. The resulting dataset was then manually modified to include the binary-coded indel data from the full plastid alignment, which was used in a partitioned mixed-data analysis (Chernomor et al., 2016) with IQ-TREE v2.3.6 (Minh et al., 2020). We performed a χ² composition test on the concatenated regions (p < 0.05; df = 1) and progressively merged partitions using ModelFinder2 (-m MF+MERGE) (Kalyaanamoorthy et al., 2017) until no further model fit improvement was observed.

We executed 100 independent maximum likelihood (ML) runs with 1,000 ultrafast bootstrap replicates (Hoang et al., 2018) to assess branch support. To reduce the risk of support overestimation, a hill-climbing nearest neighbor interchange (-bnni) search was applied. IQ-TREE was set to automatically detect the optimal number of CPU cores (-nt AUTO) to utilize the computational resources of the Puhti supercomputer at CSC, Espoo, Finland. The resulting tree was visualized using ggtree in R (Yu, 2022). All nodes with a bootstrap score below 60 were collapsed to produce a 60% consensus tree. Any node connecting two identical tips was also collapsed, but both accessions were retained in the tip-label.

### 2.6 Nuclear data assembly and analyses

We used HybPiper v2.20 (Johnson et al., 2016) to identify putative single or low-copy nuclear genes from the angiosperms353 and conserved orthologous set (COS) I and II gene sets from the high-quality raw genome skimming reads. They were mapped to the a353, COSI, and COSII (Fulton et al., 2002; Johnson et al., 2019; Wu et al., 2006) marker sets using BWA, with a coverage cutoff of 3x to maximize gene recovery. We retrieved sequences for DNA, intron, and supercontig regions for each sample. To assess assembly quality and the success of gene recovery, we used the *hybpiper stats* and the *recovery heatmap* function. We identified potential paralog sequences using the *paralog retriever* function.

### 2.7 Nuclear data preparation

Using the DNA sequences, we removed genes that gave a paralog warning by HybPiper or those that aligned to less than 30% of the samples. We combined the COSI and COSII datasets into a single COS dataset for downstream analysis. We filtered the gene alignment using BMGE (Block Mapping and Gathering with Entropy) (Galaxy Version 1.12_1) (Criscuolo & Gribaldo, 2010), which was run through the galaxy.pasteur.fr interface with default settings. We created two concatenated alignments—one for a353 and one for COS—using bad2matrix specifying -i (no indel coding), -g (no gene content coding), and -f (full FASTA names). We assessed the gene alignments for stationarity and homogeneity through a symmetry test in IQ-tree v2.3.6. Genes with a symmetry test p-value less than 0.05 were removed. We concatenated the remaining genes in bad2matrix, generating three final alignments: a353, COS, and a combined a353+COS alignment.

### 2.8 Nuclear network

We used the concatenated a353+COS alignments with all outgroups removed in Geneious to produce a phylogenetic network using the NeighborNet algorithm, a distance-based method implemented in SplitsTree. Calculations were based on uncorrected P-distances, and the network was visualized as a splits graph where splits are represented by parallel edges. To quantify the degree of tree-likeness in the resulting network, we generated delta scores for each tip and then calculated the mean delta score for each major clade.

### 2.9 Nuclear phylogeny

From each alignment, we removed all outgroups except for *S. lycopersicum,* which we retained for rooting. We inferred gene trees from the individual markers using IQ-tree with a partitioned model and 1000 bootstrap replicants. These gene trees were used to estimated coalescent based species trees for the a353, COS, and a353+COS datasets using ASTRAL v5.7.8. We specified -t 2 a quartet-based local posterior probability branch support calculation method, with *S. lycopersicum* as the outgroup.

To compare differences between the a353, COS, and a353+COS species trees, we calculated Robinson-Foulds (RF) distance (Robinson & Foulds, 1981) using IQ-Tree. We extracted alignment statistics from the filtered gene alignments as well as bootstrap support values and SH-aLRT support values from the maximum-likelihood trees. These metrics were used in the final species tree inference.

### 2.10 Tree discordance and conflict

To assess topological discordance, we used IQ-TREE to calculate gene concordance factors (gCF) by comparing each gene tree to the ASTRAL species tree. Site concordance factors (sCF) were calculated by comparing the concatenated alignments to the ASTRAL tree. These values were analyzed in R using the *Concordance* script by Lanfear (2025).

Phyparts v. 0.01 (Smith et al., 2015) was used to visualize gene tree conflict amongst the 1,000 bootstrap replicant loci trees produced previously in IQ-Tree. We first rooted all gene trees to *S. lycopersicum*, then ran Phyparts with the ASTRAL species tree as a reference and a concordance threshold of 50% (-s 0.5). We used a modified version of the phypartspiecharts script (https://github.com/mossmatters/phyloscripts) to visualize conflict or concordance of gene trees to the species tree through piecharts mapped onto the species tree nodes.

We used Phylofusion (Zhang et al., 2025), an experimental program currently in preprint, to further explore reticulate evolutionary patterns. We uploaded the trees into SplitsTree v.6.4.11(Huson & Bryant, 2024), where Phylofusion is integrated, and followed the pipeline as outlined by Huson & Bryant (2024) to prepare the files for network analyses. Then, we were able to compute rooted networks using the Phylofusion algorithm, which decomposes trees into rooted tripartitions. Reticulate edges where conflicting topologies could suggest introgression or hybridization are displayed as reticulation arches.

To assess if the observed cytonuclear conflict is reflective of ILS or chloroplast introgression, we performed a two-step test. First, we modeled the Robinson Foulds (1981) distances for 10,000 simulated trees using the contained_coalescent_tree function of DendroPy v5.0.8 (Sukumaran and Holder, 2010). We then compared the distribution of RF distances in the multispecies coalescent (MSC) simulations to the observed RF distance, with the simulations reflecting neutral cytonuclear conflict under ILS. This was done with nuclear branch lengths of 2x and 4x to model plastome coalescence. Second, we quantified how often each branch of the empirical plastome tree was recovered among simulated gene trees using the -gcf function in IQ-TREE v2.3.6. Under the null expectation of ILS, conflicting plastome branches should recur at appreciable frequency, whereas branches recovered at ≤1% were interpreted as evidence for additional processes such as chloroplast capture.

## 3. Results

### 3.1 Assembled plastome data

A total of 99 complete circularized plastid genomes were assembled from genome skimming data, representing 25 species within the Morelloid clade (Table S1 & S2). Of these, 93 plastomes were newly assembled, while the remaining were sourced from previously published, fully annotated plastomes (Table S1 & S2). For samples from which we extracted DNA, the number of recovered reads ranged from 1,718,760 to 74,540,416 (Table S2). By comparison, the Lin et al. (2022) dataset included samples with up to 95,648,594 reads (Table S1). Coverage ranged from 49 in *S. nigrum* (S29) to 1,195 in *S. triflorum* (S44), with a mean coverage of 697 (Table S1 & S2). The standard deviation for plastome coverage was 315, reflecting that the quality of DNA extractions and reads from museum specimens is variable. Plastome sizes ranged from 155,766 bp (S6, S41, S40, SP3376.1) to 154,002 bp (S8; Table S5). All plastomes retained the typical quadripartite structure, consisting of two inverted repeat (IR) regions (25,454 −25,623 bp), a large single copy (LSC) region (84,380 – 86,152 bp), and a small single copy (SSC) region (18,357 – 18,552 bp; Table S5). GC content across complete plastomes ranged from 37.9% to 38.0%, with slight variation among regions (Table S5). Most assemblies were generated using GetOrganelle, while a subset required seed sequences or assembly through NOVOPlasty to achieve circularization (Table S1 & S2). These combined strategies ensured the full recovery of the plastome across the dataset. Gene annotation identified 133 genes in each plastome. Of these genes, 89 were protein-coding, 36 were tRNAs, and 8 were rRNA (Fig. S1).

The final plastome alignment consisted of 121 sequences, totalling 104,495 bps, of which 4,305 were parsimony-informative, joined by 4,848 singletons and 95,342 constant sites. Accounting for the changes in insertion-deletion events in Morelloid plastid genomes generated an additional 1,995 binary characters, of which 1,143 were parsimony-informative while 661 represented uninformative sites.

### 3.2 Plastome phylogeny

To investigate genetic diversity within the Morelloid clade, we generated a well-supported maximum likelihood chloroplast phylogeny, applying a minimum support threshold of 60%. Nodes representing accessions with identical chloroplast sequences were collapsed into a single tip. The final tree retained 84 tips (Fig. 2). Of all the nodes in the tree, 71% received ultrafast bootstrap support of ≥95%, 23% fell between 60% and 95%, and only six nodes were below the 60% threshold and were therefore excluded (Fig. 2). Lower ultrafast values were generally limited to shallow divergences within species complexes, such as amongst *S. americanum* and *S. nigrescens* accessions, where population-level variation and potential recent divergence may reduce phylogenetic resolution.

**Fig. 2.**
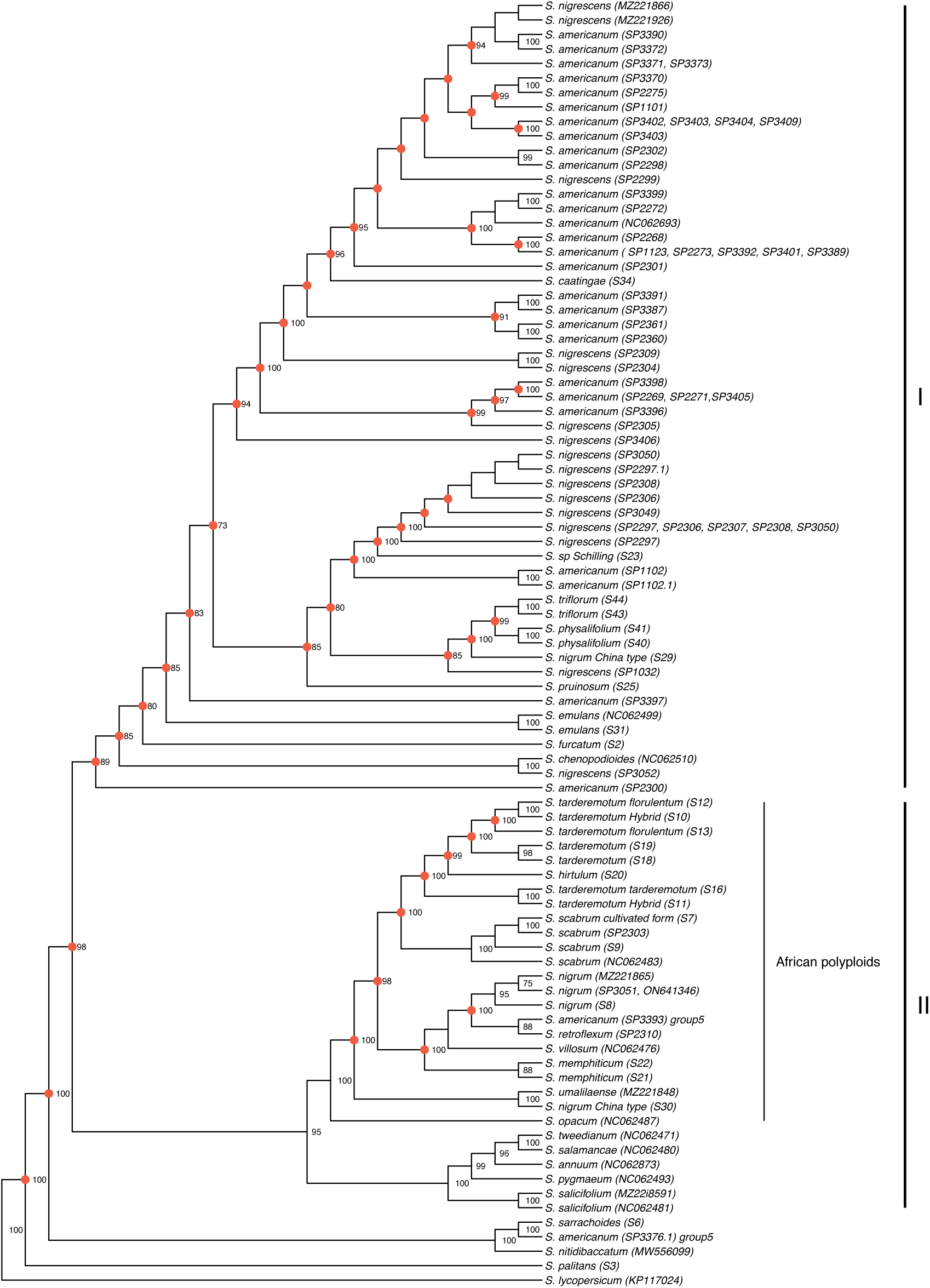
A plastome phylogeny generated from 101 complete Morelloid plastomes recovered via genome skimming. Accessions with identical chloroplast sequences were collapsed into single branches, and nodes with bootstrap support values elow 60% were excluded. Three major clades are highlighted. Branches that are affected by chloroplast capture according to MSC simulations are marked in red. Bootstrap support values are shown adjacent to each node. Newly sequenced samples are labelled with lab codes, while previously published sequences are identified by GenBank accession numbers or SRA numbers.

The resulting Morelloid chloroplast phylogeny has three major clades (Fig. 2). Clade I comprises diploid species, including *S. caatingae* and *S.* sp. Schilling sample, that are sister to Clade II, which is composed primarily of African polyploids, including *S. tarderemotum*, *S. scabrum*, and *S. nigrum* (Fig. 2; Table 2). Notably, two accessions of *S. nigrum* (China type), which exhibit morphological differences from other *S. nigrum* accessions, are resolved outside of the *S. nigrum* clade but separate from one another (Fig. 2; Table 2). Clade III, sister to Clade I and II, consists of ancestral diploid lineages, including *S. sarrachoides*, *S. nitidibaccatum*, and an accession of *S. americanum* Group 5 (Fig. 2). Finally, *S. palitan*s is found sister to the rest of the Morelloid clade (Fig. 2).

**Table 2.** Accessions with uncertain identity in the chloroplast phylogeny.

| Accession | Morphological identification | Chloroplast clade |
| --- | --- | --- |
| SP3052 (Group 5 *S. americanum*) | *S. nigrescens* | I |
| SP3376 (Group 5 *S. americanum*) | Unknown | III |
| SP2303 (Group 5 *S. americanum*) | *S. scabrum* | II (African polyploids) |
| SP2310 (Group 5 *S. americanum*) | *S. retroflexum* | II (African polyploids) |
| SP3393 (Group 5 *S. americanum*) | Unknown | II (African polyploids) |
| SP3051 (Group 5 *S. americanum*) | *S. nigrum* | II (African polyploids) |
| S29 | *S. nigrum* (China type) | I |
| S30 | *S. nigrum* (China type) | II (African polyploids) |
| S34 | *S. caatingae* | I |
| S23 | *S. sp. Schilling* | I |

Based on morphological reidentification of the Lin et al. (2022) *S. americanum* accessions, those which were originally identified as “Group 4 *S. americanum*” were reidentified as *S. nigrescens* (Table S1). All but two samples from Groups 1-3 were retained as *S. americanum* (Table S1). The Group 5 *S. americanum* accessions, which Lin et al. (2022) expected to be misidentified, are found in clades I, II, and III, with four of them in the African polyploids (Table 2).

Out of 118 plastome nodes, 47 (∼40%) showed <1% recovery in simulated gene trees – far below the frequency expected under incomplete lineage sorting (ILS). These low-frequency conflicting plastome branches were found primarily in Clades I and II (highlighted in red, Fig. 2) and are more consistent with historical chloroplast introgression than with neutral ILS. Most remaining discordant nodes occurred at appreciable frequencies (>1%) in simulations, consistent with ILS as a plausible source of those topology incongruences.

### 3.5 Nuclear gene recovery and final alignments

HybPiper successfully recovered 349 genes from the a353 marker set, 321 genes from the COSII marker set, and 1,807 genes from the COSI marker set (Table S4). Genes were removed from the analysis if found in fewer than 30% of samples, gave paralog warnings, or failed the symmetry test (Table S6). As a result, 296 a353 genes, 4 COSI genes, and 107 COSII genes were retained for downstream analyses. Four samples (S9, S31, ERR1042995, and ERR10429955) were removed from analysis due to low overall recovery rates. The COSI and COSII genes were merged into a single dataset.

Three alignments were generated based on the a353 genes, the COS genes, and a combination of the two (Table 3). The a353 alignment is longer than the COS alignment and has a high proportion of constant sites (0.88; Table 3). The COS alignment has a greater overall number of parsimony-informative sites (Table 3). Based on the maximum-likelihood trees produced from these alignments, the combined a353 and COS tree is the most informative, with the highest mean and the lowest standard deviation for both bootstrap and SH-aLRT support values (Table 3). Robinson-Foulds (RF) distances were used to compare the phylogenetic trees derived from different marker sets. The COS tree was most distinct from the a353 and the combined COS/a353 tree, with an RF distance of 350 for both comparisons. The RF distance between the a353 tree and the combined tree (114) suggests that adding COS markers does not drastically alter the a353 tree. As a result, the combined tree was used for the final nuclear phylogeny.

**Table 3.** Alignment and phylogenetic support statistics from maximum-likelihood gene tree analyses.

|  | **a353** | **COS** | **a353+COS** |
| --- | --- | --- | --- |
| **Length** | 243,054 bp | 59,228 | 302,282 |
| **Parsimony-informative sites** | 1,904 bp | 3,836 bp | 5,740 |
| **# Constant sites** | 214,969 | 44,066 | 259,035 |
| **Mean bootstrap** | 69.9 | 77.1 | 77.4 |
| **StDev bootstrap** | 24.7 | 18.3 | 17.5 |
| **Mean SH-aLRT** | 83.6 | 84.2 | 87.5 |
| **StDev SH-aLRT** | 23.4 | 23.7 | 16.3 |

### 3.7 Nuclear phylogeny

For nuclear phylogeny, sCF and gCF were generated for each node. Concordance factors cluster between 0 and 25 for gCF and between 25 and 50 for sCF (Fig. 3), indicating that overall, individual sites strongly support a branch, but there is more conflict amongst gene trees, likely a sign of incomplete lineage sorting or hybridization. Higher bootstrap values are associated with higher gCF values, suggesting that these branches are better supported (Fig. 3).

**Fig. 3.**
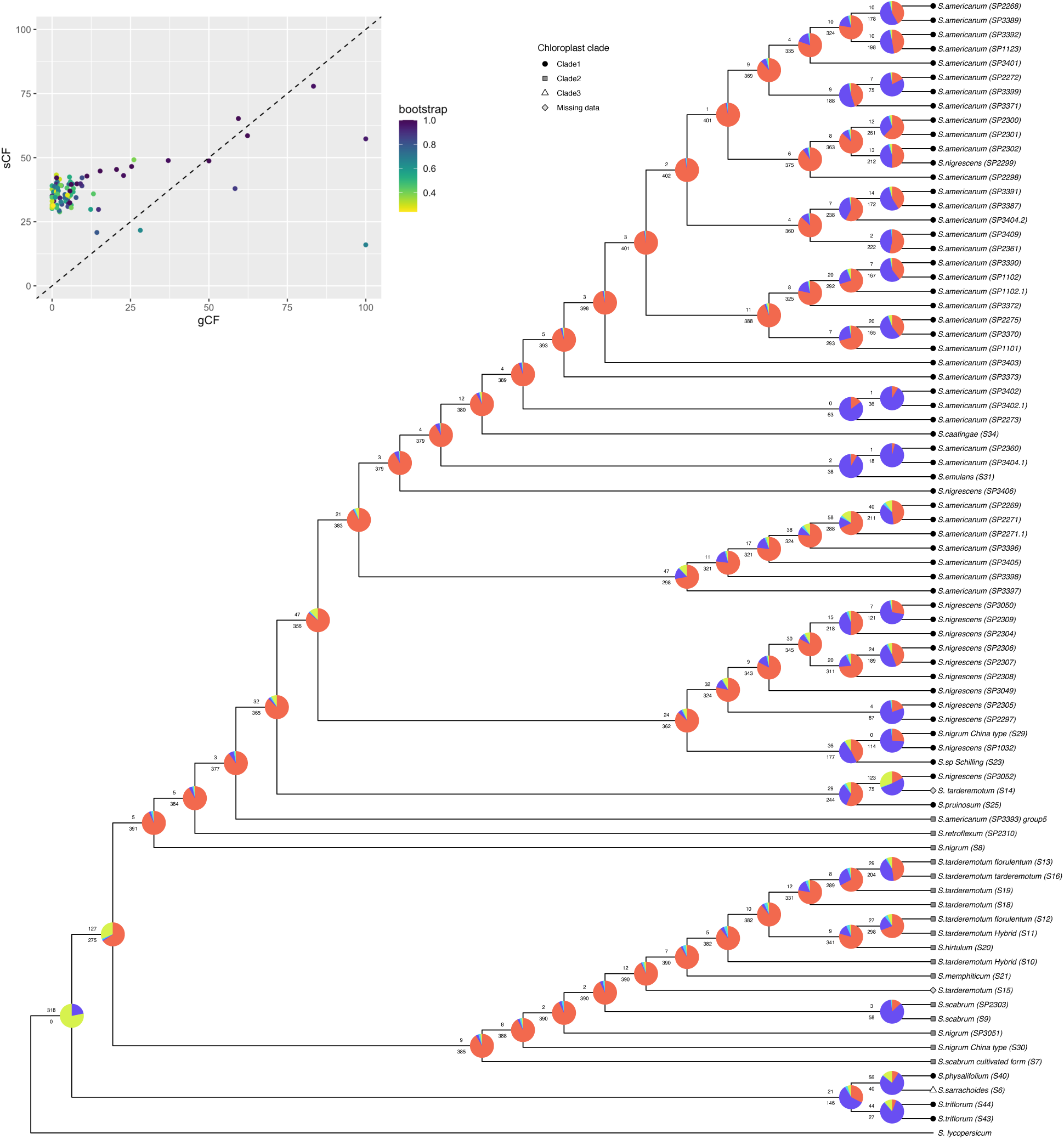
Nuclear phylogeny with concordance factors. Panel A shows the nuclear phylogeny, including the phyparts pie charts where turquoise represents the top alternative bipartition, red represents all other alternative bipartitions, blue represents uninformative bipartitions for that node, and green represents gene trees in concordance. The number at the top of each branch indicates the number of concordant gene trees, and the lower number indicates conflicting gene trees. In the upper panel, concordance factors are based on a combined set of a353 and COS markers from genome skimming, incorporating bootstrap values as a color gradient. gCF: The proportion of gene trees that support a particular branch in the species tree. sCF: The proportion of sites in the sequence alignment that support a particular branch. The dashed diagonal line represents the gCF = sCF relationship, where points above the line indicate higher site support than gene tree support, while points below the line suggest higher gene tree support relative to site-based support.

The proportion of concordant to conflicting gene trees for a given node is the highest for the split between Clade I and Clade II, with 127 of 275 gene trees concordant (Fig. 3). Clade III also has a relatively higher proportion, although many gene trees are uninformative (Fig. 3). Beyond this, there is low concordance with multiple nodes having fewer than five gene trees supporting the given relationship.

The nuclear network splits the Morelloid clade into six groups (Fig. 4). Groups A and F are made up of *S. americanum* samples except for an *S. nigrescens* in Group F (Fig. 4). Group B includes all other *S. nigrescens* samples, an *S. nigrum* China type, and the Schilling sample. Groups C, D, and E are largely made up of the polyploid accessions; Group D includes all but one of the *S. tarderemotum* accessions (Fig. 4). Group A is preceded by a characteristic split and is clearly distinct from the rest (Fig. 4).

**Fig. 4.**
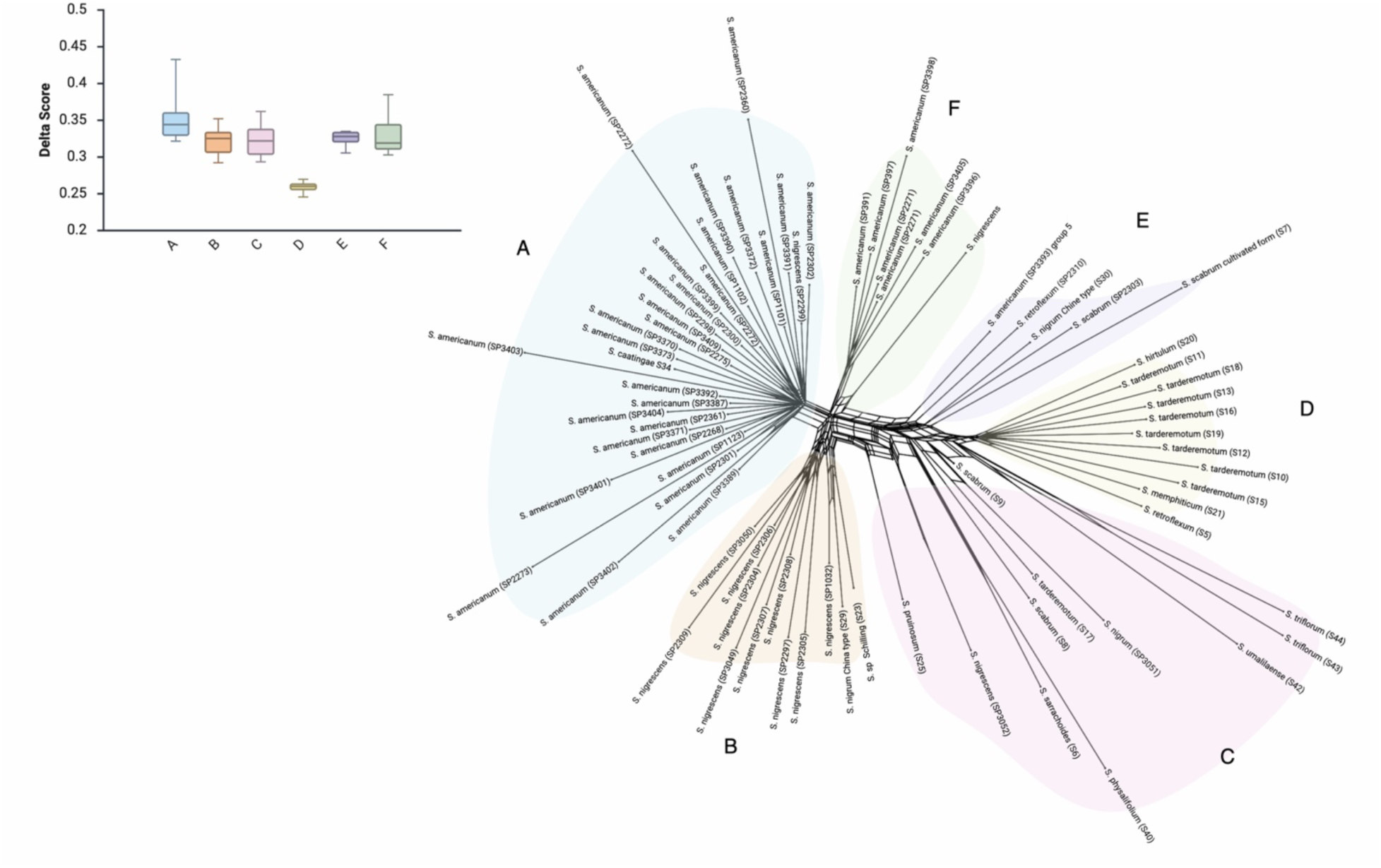
Nuclear phylogenetic network produced from the combination of a353 and COS dataset and calculated using the NeighborNet algorithm. Major clades of the network are highlighted with the color corresponding to the delta scores box plots. Clades in are labeled A through F, starting at the far-left clades and continuing counterclockwise.

Delta scores ranged from 0.3 to 0.4, indicating a moderate degree of tree likeness (Fig. 4). The greatest variation in delta scores, as well as the highest values, is in Clade I (Fig. 4). Conversely, Clade IV shows the least variation and the lowest delta scores, indicating it is the least tree-like (Fig. 4).

### 3.8 Cytonuclear discordance

The simulation with 4× scaling of nuclear branch lengths produced a null distribution of RF distances ranging from around 135 to below 147.5, with the peak centered at ∼145 and a frequency above 3,500 (Fig. S2). The observed RF distance between the empirical plastome tree and nuclear tree is 130 with a P-value of 1.000. This is significantly below what would be expected from ILS alone, as observed in the simulations. Several clades exhibited pronounced discordance relative to the nuclear species tree, indicating localized cytonuclear incongruence consistent with historical reticulation (Fig. 5). In these cases, plastomes are phylogenetically associated with lineages that are clearly distinct in the nuclear analysis, suggesting past hybridization followed by backcrossing and the introgression of plastomes across species boundaries while retaining divergent nuclear genomes. Extensive reticulation is apparent amongst the Morelloids, particularly in the *S. americanum* group (Fig. 5). This is true not only within *S. americanum* and *S. nigrescens* but also between the two, indicating substantial admixture in these populations (see also Lin et al., 2022). Simulations confirmed that such mixed groupings are virtually never produced by ILS alone (recovered in ≤1% of gene trees), implying repeated chloroplast exchange between these lineages. Some shallow-level conflicts among the African polyploids occurred more frequently under the coalescent model, suggesting that recent rapid radiations can yield alternative gene-tree resolutions without requiring introgressive chloroplast transfer. Much of these reticulations are within the *S. tarderemotum* complex and between the *S. tarderemotum*, *S. scabrum*, and *S. nigrum* accessions (Fig. 5). Two reticulate arches connect the *S. americanum* accessions to the polyploids (Fig. 5).

**Fig. 5.**
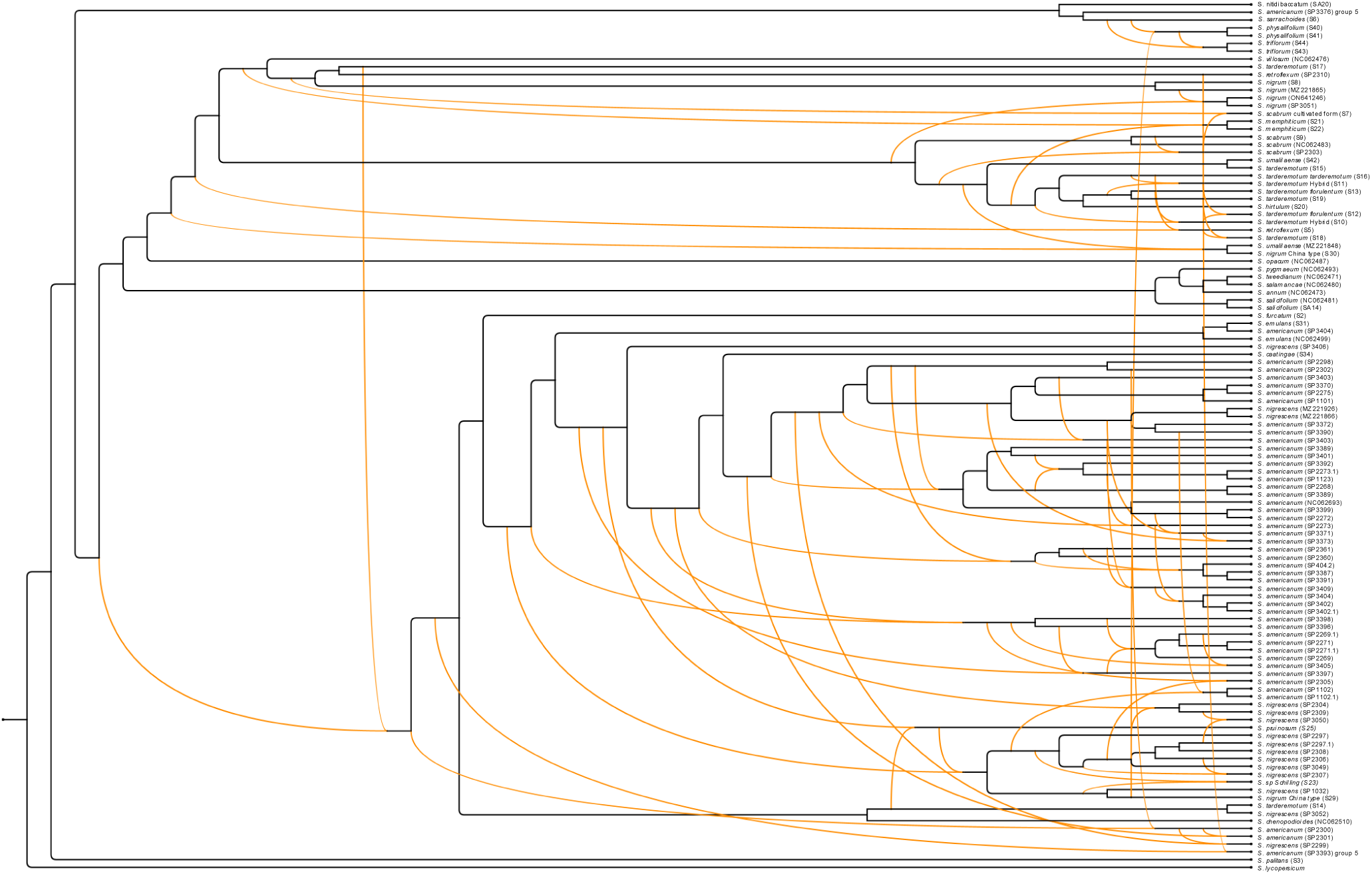
Fused rooted nuclear chloroplast tree. Orange lines represent reticulations arising from conflicting topologies. Accessions that were not included in the nuclear tree are present here, but reticulation cannot be assessed. di

## 4. Discussion

### 4.1 Evidence of reticulate evolution

Here, we present the first application of genome skimming to elucidate the evolutionary relationships within the Morelloid clade of Solanum using complete plastid genomes and a large set of nuclear genes. Past studies of the Morelloid clade of *Solanum* have relied on marker-based methods to construct chloroplast and nuclear phylogenies (Bohs, 2005; Manoko, 2024; Poczai & Hyvönen, 2011; Särkinen et al., 2015). While these approaches allow focused investigation of phylogenetically informative genes, they often suffer from limited resolution and gene-tree incongruence.

Our nuclear tree hints to reticulate evolution through conflict-dominant topologies in gene trees, low gene concordance factors, and moderate delta scores (Fig. 3). The inclusion of the chloroplast phylogeny reveals widespread cytonuclear discordance throughout the Morelloid clade, highlighted by the prevalence of reticulate arches in the fused nuclear chloroplast tree (Fig. 5). These patterns are most consistent with historical introgression events involving chloroplast exchange, particularly within the pan-American diploids (Clade I) and African polyploids. Below, we discuss examples of discordance and potential evolutionary histories throughout the Morelloid clade.

### 4.2 Morelloid diploid lineages

In the plastome phylogeny, *S. americanum* and *S. nigrescens* are nested within Clade I (Fig. 2). The nuclear phylogeny resolves all *S. americanum* samples in a single clade and *S. nigrescens* samples in a closely related one (Fig. 3). There is pervasive reticulation between the *S. americanum* and *S. nigrescens* accessions, suggesting that the lineages have undergone introgression, perhaps reflecting the broad tropical distribution of these weeds (Fig. 5). Based on MSC simulations, the observed patterns are more consistent with chloroplast capture than with ILS, suggesting the plausibility of historical introgression facilitating plastid exchange.

As such, the widespread Pan-American diploids form a well-supported clade I made up primarily of *S. americanum* and *S. nigrescens*. Notably, *S. americanum* is a pantropical weed, and its ecology suggests it could have hybridized opportunistically. For example, there are herbarium specimens in Tanzania identified as *Solanum* aff. *americanum* or *S. nigrum,* which were ultimately determined to be other species (Särkinen et al., 2018). Despite this, neither *S. americanum* nor *S. nigrescens* is a direct ancestor of the African polyploids. They did not contribute chloroplasts to the Afro-tropical polyploids we sampled and instead are confirmed as a distinct Pan-American lineage.

*Solanum nitidibaccatum*, *S. sarrachoides*, and *S. palitans* are the earliest branching species in the plastome phylogeny with 100% bootstrap support (Clade III), suggesting their role as ancestral diploid lineages within the Morelloid clade (Fig. 2). We were unable to include *S. palitans* or *S. nitidibaccatum* in the nuclear phylogeny and, therefore, may only draw conclusions on their plastome contribution. In the nuclear phylogeny, *S. physalifolium* and *S. triflorum* group with *S. sarrachoides* at the early branching positions of the tree (Fig. 2). *Solanum triflorum* and *S. physalifolium* group with the Pan-American diploids (Clade I) in the chloroplast phylogeny (Fig. 2). As such, these species’ nuclear lineage was an early-diverging member of the Morelloids, but their maternal plastid lineage is not an early contributor; instead, it resembles the plastid types of *S. nigrescens*.

The South American *S. tweedieanum*, *S. salamancae*, *S. annuum*, *S. pygmaeum*, and *S. salicifolium* form a well-supported clade in the chloroplast phylogeny that shares a most recent common ancestor with the African polyploids (Fig. 2). Among these, *S. tweedieanum*, *S. pygmaeum*, and *S. salicifolium* are classified as diploids based on chromosome counts, but no cytometric data are available (Särkinen et al., 2015). The ploidy of *S. annuum* and *S. salamancae* is unknown. It is unexpected that these species group more closely with polyploid species than with the diploid species of Clades I and III. These patterns could reflect processes such as chloroplast capture, diploidization following an earlier polyploidization event or other sources of cytonuclear discordance. Within the *Solanum* genus, autopolyploidization appears to have occurred repeatedly, as ten potato autotetraploids were observed in a study by Hardigan et al. (2017). This pattern in closely related species supports the hypothesis that species like *S. annuum* or *S. salamancae* may be ancient autotetraploids that have undergone sufficient diploidization to obscure their genomic history. However, nuclear data is necessary to test whether these species are indeed diploids that contributed to the African polyploid plastid type.

### 4.3 Diploid contributions to African polyploids

The African polyploid plastome lineage forms a distinct monophyletic clade (Fig. 2). No Pan-American diploid plastomes (Clade I) show introgression, suggesting that these lineages originated in Africa or were derived from Afro-Eurasian ancestors. One Chinese accession of *S. nigrum* (China-type S30) even carries this African plastome, pointing to dispersal or secondary contact beyond Africa. This clade is nested among diploid species lineages, and one or more of these appear to have donated the plastids that gave rise to the African plastome. In the nuclear phylogeny, *S. retroflexum* and *S. nigrum* samples are embedded amongst the Pan-American lineages (Fig. 3). As a widespread circumtropical weed, *S. americanum* may have contributed to African diversity on the paternal side. The distinct African plastome could be consistent with one or more chloroplast capture events, whereas nuclear patterns may suggest paternal contribution from *S. americanum*. These observations together are consistent with a possible hybrid origin of the polyploid clade, although additional genomic analysis would be needed to more rigorously test this hypothesis.

### 4.4 The evolution of African polyploids

*Solanum scabrum* and *S. nigrum* are both Afro-Eurasian hexaploid species introduced to the Americas through human activities. In this study, we included both wild and cultivated forms of *S. scabrum*, which, in the plastome phylogeny, form a single cluster sister to the *S. tarderemotum* complex (Fig. 2). Similarly, *S. nigrum* samples form a cluster, sister to SP3393 and SP2310, which are two *S. americanum* Group 5 accessions identified by Lin et al. (2022). Although SP2310 is reclassified as *S. retroflexum*, these accessions share plastomes with the *S. nigrum* accessions (Fig. 2). In the nuclear phylogeny, the relationship between *S. scabrum* and *S. nigrum* remains, with two *S. scabrum* clustering together as sister to two *S. nigrum* accessions, one of which is the Chinese form (Fig. 3). This, in turn, is sister to the cultivated form of *S. scabrum* (Fig. 2). *Solanum scabrum* and *S. nigrum* have prevalent reticulate arches between and amongst the complexes, as well as with *S. tarderemotum*. These patterns may be consistent with introgression and potential chloroplast capture, similar to those inferred among the Pan-American lineages (Fig. 5).

The inclusion of two “*S. nigrum*” accessions from China has revealed unexpected diversity. In both phylogenies, one Chinese sample (China-type S29) falls within the Pan-American clade, and the other (China-type S30) falls within the African hexaploids (Fig. 2 and 3). Morphological differences between Asian and European *S. nigrum* accessions have been described but are considered to be part of a continuous variation observed within *S. nigrum* across Eurasia (Särkinen et al., 2018). Notably, the Chinese *S. nigrum* has been recognised as a distinct taxon in the past, described as *S. ganchouenense* H.Lév. and *S. chenopodiifolium* H.Lév. The fact that the Chinese *S. nigrum* accessions do not appear to be closely related to our other *S. nigrum* accessions suggests the presence of cryptic diversity within the species, though further sampling is needed to confirm the presence of distinct Chinese species.

Based on their nuclear SNP phylogeny, Lin et al. (2022) expected SP2303, SP2310, SP3393, and SP3051 to be polyploids likely misidentified as *S. americanum*. This is confirmed by our nuclear and chloroplast data as well as morphological re-identification (Fig. 2 and 3; Table S1).

Interestingly, these Group 5 polyploids all differ in morphological identity and are inconsistent in their placement in the nuclear and chloroplast phylogenies. For example, SP2310 and SP3393 are closely related in both nuclear and chloroplast phylogenies. In the nuclear phylogeny, they are sister to *S. nigrum* (S8), although in a clade largely composed of diploids, and in the chloroplast phylogeny, they are sister to the *S. nigrum* accessions. *Solanum nigrum* (SP3051) is between *S. scabrum* (S9) and *S. nigrum* China type (S30) in the nuclear phylogeny. Its chloroplast is identical to that of *S. nigrum* accession ON641346. This inconsistent placement suggests incomplete genomic stabilization and mixed parental contributions. Polyploidy occurs frequently in *Solanum,* but only a few species stabilize into established species boundaries.

Based on nuclear and chloroplast phylogenies, these samples exhibit a mixture of parental genomes and polyploid identities, suggesting these accessions represent polyploids that formed but did not persist. They are likely “unstable” or “ephemeral” polyploids.

Past work has described unclear species boundaries and potential interspecific hybridization within the *S. tarderemotum* complex (Manoko, 2024; Olet, 2004; Särkinen et al., 2018). Our results show a deep web of reticulation in the entire clade apparent in the fused rooted nuclear chloroplast tree, where *S. hirtulum* is linked to the *S. tarderemotum* accessions by a reticulate arc (Fig. 5). This supports a hybrid, allopolyploid origin for *S. tarderemotum*. *Solanum hirtulum* is embedded in the *S. tarderemotum* complex in both the nuclear and plastome phylogenies, indicating that they share a recent ancestor (Figs. 2 and 3). The nuclear phylogenetic network indicates that this clade arose from a rapid radiation event and is the least tree-like clade (Fig. 4). In the nuclear phylogeny, *S. memphiticum* is also present in the *S. tarderemotum* clade, suggesting it is a contributor to the paternal lineage (Fig. 3). Formal tests confirm that the observed discordance between organellar and nuclear phylogenies is consistent with chloroplast capture, in which some *S. tarderemotum* accessions appearing to have inherited their plastome from a *S. hirtulum*-like maternal lineage (Fig. 2 and 5). One possible interpretation is that *S. tarderemotum* represents an allotetraploid hybrid, involving a *S. hirtulum*-type maternal lineage and a *S. memphiticum*-type paternal lineage. However, additional sampling and confirmation of ploidy in *S. hirtulum* would be necessary to test this hypothesis. This interpretation aligns with morpho-cytological data: *S. tarderemotum* has deeply stellate corollas, unlike *S. memphiticum*.

Earlier crossing experiments reported crosses between *S. tarderemotum* and *S. memphiticum* (Olet et al., 2015). This hypothesis also illustrates how allopolyploids can perpetuate cytonuclear conflict: nuclear markers group S. *tarderemotum* with one parent, while organelles trace to the other.

### 4.5 Evolution of Black Nightshades

Introgression is a major evolutionary driver in the Morelloids. The extent of cytonuclear discordance in our dataset appears greater than expected under neutral incomplete lineage sorting alone and is consistent with repeated chloroplast capture. Similar evolutionary patterns have been observed in Rosaceae, which includes many polyploid taxa and where reticulate evolution and hybrid origins are observed in clades such as *Potentilla* and Maleae (Hodel et al., 2022; Persson et al., 2020). As well as in *Vitis*, where studies reveal widespread cytonuclear conflict and extensive reticulate evolution (Nie et al., 2023; Talavera et al., 2023; Zecca et al., 2020). In the case of the Morelloids, our phylogenomic synthesis, combining plastome and nuclear trees, ILS tests, and known ploidy patterns, reveals a web of cytonuclear discordance resulting from widespread chloroplast capture.

## 5. Conclusions

Our results suggest that the Morelloids originated in the Neotropics with ancestral lineages including *S. palitans* and *S. sarrachoides*. The circumtropical weed, *S. americanum*, as well as the Pan-American *S. nigrescens*, represent the early diverging diploid lineages. The Morelloid clade of Solanum is well-supported, but within it, reticulation and particularly chloroplast capture, are pervasive. The African continent hosts several independently derived polyploid clades, including the *S. scabrum* and *S. tarderemotum* complexes. These African polyploids are sister to the Pan-American diploids, where chloroplast capture events may have contributed to the African diversity. The evolution of Morelloids in Africa has been driven by widespread allopolyploidy and introgression, with plastome diversity reflecting a network of transfers. This reticulation aligns with classical studies, which have noted that species boundaries are blurred by polyploidy and hybridization (Edmonds, 1977, 1978).

## CRediT authorship contribution statement

**Sundre Winslow:** Conceptualization, Data curation, Formal analysis, Funding acquisition, Visualization, Writing – original draft **Sandra Knapp**: Formal analysis, Writing – review and editing **Tiina Särkinen**: Conceptualization, Data curation, Supervision, Writing – review and editing **Péter Poczai**: Conceptualization, Data curation, Formal analysis, Funding acquisition, Supervision, Visualization, Writing – review and editing

## Acknowledgements

This work was supported by the Societas pro Fauna et Flora Fennica Master’s Thesis Grant, EU Grant ID: 101151612, and TFK Grant ID: 6306124. We thank the curators and staff of the LUOMUS Botanical Collections (HEL), and the Herbarium of the Royal Botanic Garden Edinburgh (E) for access to herbarium material. We are also grateful for the comments by Rocío Deanna on an earlier version of the manuscript. The authors wish to acknowledge CSC – IT Center for Science, Finland, for computational resources. We thank Elina Laiho for assistance from the Historical DNA Laboratory. We are grateful to the authors of previously published datasets, particularly Lin et al. (2022) and Gagnon et al. (2022), for making their genomic data publicly available.

