## Supplemental Table 1-3 and Supplemental Figures for "Cytonuclear Conflict and Reticulate Evolution in the Morelloid Clade (*Solanum*, Solanaceae): Insights from Genome Skimming and Network Phylogenomics"

**Supplementary Table 1: Samples used from Lin et al. (2022) with reference to the group number the accession fell in their phylogeny.**

| Accession | Group | Species | Barcode | Assembly method | No. Reads | Plastome coverage |
| --- | --- | --- | --- | --- | --- | --- |
| SP3409 | 3 | *S. americanum* | BM014636780 | GetOrganelle | 68492244 | 1013.11 |
| SP3406.1 | 1 | *S. nigrescens* | BM014636772 | GetOrganelle | 18356480 | 786.45 |
| SP3405.1 | 1 | *S. americanum* | None | GetOrganelle | 15370814 | 854.05 |
| SP3404.1 | 3 | *S. americanum* | BM014636767 | GetOrganelle | 13457210 | 556.11 |
| SP3403.1 | 3 | *S. americanum* | BM014636774 | GetOrganelle | 17466000 | 864.75 |
| SP3402.1 | 3 | *S. americanum* | BM014636849 | GetOrganelle | 31717608 | 887.88 |
| SP3401.1 | 3 | *S. americanum* | BM014636824 | GetOrganelle | 16751914 | 933.18 |
| SP3399.1 | 3 | *S. americanum* | BM014636832 | GetOrganelle | 13390954 | 922.57 |
| SP3398.1 | 1 | *S. americanum* | BM014636786 | GetOrganelle | 20661176 | 962.74 |
| SP3397.1 | 1 | *S. americanum* | BM014636828 | GetOrganelle | 18748284 | 879.14 |
| SP3396.1 | 1 | *S. americanum* | BM014636789 | GetOrganelle | 10996382 | 822.96 |
| SP3393 | 5 | *S. americanum* | None | GetOrganelle | 71834388 | 908.94 |
| SP3392 | 3 | *S. americanum* | BM014636831 | GetOrganelle | 67990370 | 1014.6 |
| SP3391.1 | 3 | *S. americanum* | BM014636825 | GetOrganelle | 12383492 | 803.21 |
| SP3390.1 | 2 | *S. americanum* | BM014636853 | GetOrganelle | 13943028 | 1043.44 |
| SP3389 | 3 | *S. americanum* | BM014636775 | GetOrganelle | 75199924 | 641.54 |
| SP3387 | 3 | *S. americanum* | BM014636765 | GetOrganelle | 72375748 | 826.15 |
| SP3376 | 5 | *S. americanum* | None | GetOrganelle | 24032913 | 954.42 |
| SP3373.1 | 3 | *S. americanum* | None | GetOrganelle | 7905646 | 927.31 |
| SP3372.1 | 2 | *S. americanum* | BM014636785 | GetOrganelle | 15425370 | 886.07 |
| SP3371.1 | 3 | *S. americanum* | BM014636852 | GetOrganelle | 7555198 | 627 |
| SP3370.1 | 2 | *S. americanum* | BM014636843 | GetOrganelle | 11911262 | 809.29 |
| SP3052 | 5 | *S. nigrescens* | BM014636845 | GetOrganelle | 69827018 | 916.52 |
| SP3051 | 5 | *S. nigrum* | BM014636835 | GetOrganelle | 21415932 | 842.84 |
| SP3050 | 4 | *S. nigrescens* | BM014636840 | GetOrganelle | 68041404 | 912.16 |
| SP3049.1 | 4 | *S. nigrescens* | BM014636837 | GetOrganelle | 12286168 | 782.12 |
| SP2361 | 3 | *S. americanum* | BM014636846 | GetOrganelle | 76427690 | 782.51 |
| SP2360.1 | 3 | *S. americanum* | BM014636833 | GetOrganelle | 26116316 | 872.39 |
| SP2310 | 5 | *S. retroflexum* | BM014636836 | GetOrganelle | 83637218 | 812.19 |
| SP2309 | 4 | *S. nigrescens* | BM014636777 | GetOrganelle | 9585248 | 1005.54 |
| SP2308 | 4 | *S. nigrescens* | BM014636839 | GetOrganelle | 74834512 | 748.51 |
| SP2307.1 | 4 | *S. nigrescens* | BM014636773 | GetOrganelle | 9585248 | 898.85 |
| SP2306 | 4 | *S. nigrescens* | BM014636841 | GetOrganelle | 95648594 | 825.95 |
| SP2305.1 | 4 | *S. nigrescens* | BM014636788 | GetOrganelle | 12760232 | 845.63 |
| SP2304 | 4 | *S. nigrescens* | BM014636778 | GetOrganelle | 66657676 | 956.77 |
| SP2303 | 5 | *S. scabrum* | BM014636827 | GetOrganelle | 60357468 | 798.16 |
| SP2302 | 3 | *S. americanum* | BM014636848 | GetOrganelle | 74440626 | 922.68 |
| SP2301 | 3 | *S. americanum* | BM014636779 | GetOrganelle | 73283534 | 817.29 |
| SP2300 | 3 | *S. americanum* | BM014636850 | GetOrganelle | 71417766 | 789.81 |
| SP2299 | 3 | *S. nigrescens* | BM014636847 | GetOrganelle | 71495954 | 867.51 |
| SP2298.1 | 3 | *S. americanum* | BM014636784 | GetOrganelle | 10964434 | 951.91 |
| SP2297.1 | 4 | *S. nigrescens* | BM014636764 | GetOrganelle | 16338266 | 794.49 |
| SP2275 | 2 | *S. americanum* | BM014636783 | GetOrganelle | 66343806 | 838.13 |
| SP2273.1 | 3 | *S. americanum* | BM014636781 | GetOrganelle | 46318972 | 781.18 |
| SP2272.1 | 3 | *S. americanum* | BM014636851 | GetOrganelle | 16283590 | 907.45 |
| SP2271.1 | 1 | *S. americanum* | BM014636776 | GetOrganelle | 72608494 | 923.32 |
| SP2269.1 | 1 | *S. americanum* | None | GetOrganelle | 12664336 | 923.21 |
| SP2269.2 | 1 | *S. americanum* | None | GetOrganelle | 55611512 | 923.21 |
| SP2268.1 | 3 | *S. americanum* | BM014636842 | GetOrganelle | 57947206 | 847.83 |
| SP1123 | 2 | *S. americanum* | BM014636830 | GetOrganelle | 73550082 | 948.06 |
| SP1102.2 | 2 | *S. americanum* | BM014636790 | GetOrganelle | 64872174 | 814.55 |
| SP1101 | 2 | *S. americanum* | BM014636844 | GetOrganelle | 71087646 | 842.44 |
| SP1032 | 4 | *S. nigrescens* | BM014636834 | GetOrganelle | 36674439 | 827.46 |

Supplementary Table 2: Newly sequenced samples

| Lab code | Accession | Species | Barcode | Herbarium | Assembly methods | No. Reads | Coverage | Locality |
| --- | --- | --- | --- | --- | --- | --- | --- | --- |
| S10 | PV802119 | *S. tarderemotum* | A14750151 | Nijmegen | GetOrganelle | 50915806 | 831.56 | Kenya |
| S11 | PV802118 | *S. tarderemotum* | A14750182 | Nijmegen | NOVOPLASTY | 44914572 | 833.83 | Kenya |
| S12 | PV802117 | *S. tarderemotum* | A14750164 | Nijmegen | NOVOPLASTY | 43169001 | 304.01 | Kenya: Kari-Kisii |
| S13 | PV802116 | *S. tarderemotum* | A14750173 | Nijmegen | GetOrganelle | 68874162 | 326.31 | Kenya: Londiani |
| S16 | PV802115 | *S. tarderemotum* | A34750041 | Nijmegen | GetOrganelle | 35996701 | 198.95 | Kenya: Migori district |
| S18 | PV802114 | *S. tarderemotum* | A34750047 | Nijmegen | GetOrganelle | 29018647 | 236.27 | Kenua: Kisii district |
| S19 | PV802113 | *S. tarderemotum* | A34750047 | Nijmegen | NOVOPLASTY | 34969358 | 167.53 | Kenya: Kisii district |
| S2 | PV802134 | *S. furcatum* | C.32173 | Helsinki | GENEIOUS | 54471856 | 389.04 | Chile |
| S20 | PV802134 | *S. hirtulum* | E00526867 | RBGE | GetOrganelle | 68260988 | 465.92 | Ethiopia |
| S21 | PV802133 | *S. memphiticum* | E00621066 | RBGE | NOVOPLASTY | 39423939 | 942.63 | Saudi Arabia |
| S22 | PV802132 | *S. memphiticum* | E00621077 | RBGE | NOVOPLASTY | 63356308 | 909.58 | Yemen |
| S23 | PV802120 | *Schilling* |  |  | GetOrganelle | 42382079 | 852.66 | USA |
| S25 | PV802126 | *S. pruinosum* | MEXU:1207806 | MEXU | GetOrganelle | 61180420 | 526.89 | Mexico |
| S29 | PV802131 | *S. nigrum (china)* | E00133328 | RBGE | NOVOPLASTY | 40634775 | 49.86 | China |
| S3 | PV802129 | *S. palitans* | C.322687 | Helsinki | NOVOPLASTY | 44838349 | 1005.84 | Bolivia |
| S30 | PV802130 | *S. nigrum (china)* | E00320320 | RBGE | NOVOPLASTY | 38559958 | 864.46 | China |
| S31 | PV802135 | *S. emulans* | E00526730 | RBGE | GetOrganelle | 10753687 | 981.15 | USA |
| S34 | PV802136 | *S. caatingae* | RB01145303 | RBGE | NOVOPLASTY | 37884182 | 875.43 | Brazil, Paraíba, Carrapateira, |
| S40 | PV802128 | *S. physalifolium* | 954750160 | Nijmegen | GetOrganelle | 33347642 | 227.07 |  |
| S41 | PV802127 | *S. physalifolium* | 964750073 | Nijmegen | GetOrganelle | 28356314 | 987.07 |  |
| S43 | PV802112 | *S. triflorum* | C.322930 | Helsinki | GetOrganelle | 26958882 | 1092.73 | Alberta, Canada |
| S44 | PV802111 | *S. triflorum* | C.322927 | Helsinki | GetOrganelle | 26249349 | 1195.28 | Saskatchewan, Canada |
| S6 | PV802124 | *S. sarrachoides* | A44750069 | Nijmegen | GetOrganelle | 72427348 | 893.77 | none |
| S7 | PV802123 | *S. scabrum cultivated* | C.322816 | Helsinki | GetOrganelle | 59345692 | 199.34 |  |
| S8 | PV802122 | *S. scabrum wild* | C.322809 | Helsinki | NOVOPLASTY | 33304282 | 893.48 | Sudan |
| S9 | PV802121 | *S. scabrum* | 994750011 | Nijmegen | GetOrganelle | 57002260 | 810.42 | Foumbot, Cameroon |

Supplementary Table 3: **Previously Published Fully Annotated Plastomes Used in This Study**

| Accession | Species | Voucher | Locality |
| --- | --- | --- | --- |
| ON641346 | *S. nigrum* | LISE:96441 | Portugal |
| NC062693 | *S. americanum* | none |  |
| NC062481 | *S. salicifolium* | HFN:s.n. |  |
| NC062471 | *S. tweedianum* | CORD:Barboza 2229 |  |
| MZ221866 | *S. nigrescens* | HFN:A14750130 |  |
| MZ221926 | *S. nigrescens* | BM:Craat et al 97419 |  |
| MZ221848 | *S. umalilaense* | HFN:A14750133 |  |
| MZ221865 | *S. nigrum* | NY:Nee 45551 |  |
| NC062476 | *S. villosum* | A14750161a (NIJ) | Kenya: Maseno |
| NC062483 | *S. scabrum* | Nee 45551 (NY) | Australia: Adelaide |
| NC062487 | *S. opacum* | Bohs 3561 (UT) | Auatralia: Queensland |
| NC062873 | *S. annum* | Barboza 3017 (CORD) | Argentina: Tucuman |
| NC052480 | *S. salamancae* | Barboza 2182 (CORD) | Argentina: Tucuman |
| NC062493 | *S. pygmaeum* | King 301 (BM) | none |
| MZ221859 | *S. salicifolium* | Knapp 10492 (BM) | Argentina, Mendoza, San Carlos |
| NC062510 | *S. chenopodioides* | Barboza et al. 2316 (CORD) | Argentina: Buenos Aires |
| NC062499 | *S. emulans* | NYBG1cult. | USA, NY |
| MW556099 | *S. nitidibaccatum* | Chiarini et al. 795 (CORD) | Argentina; Cordoba |
| NC062471 | *S. tweedianum* | Barboza 2229 | Kenya: Maseno |

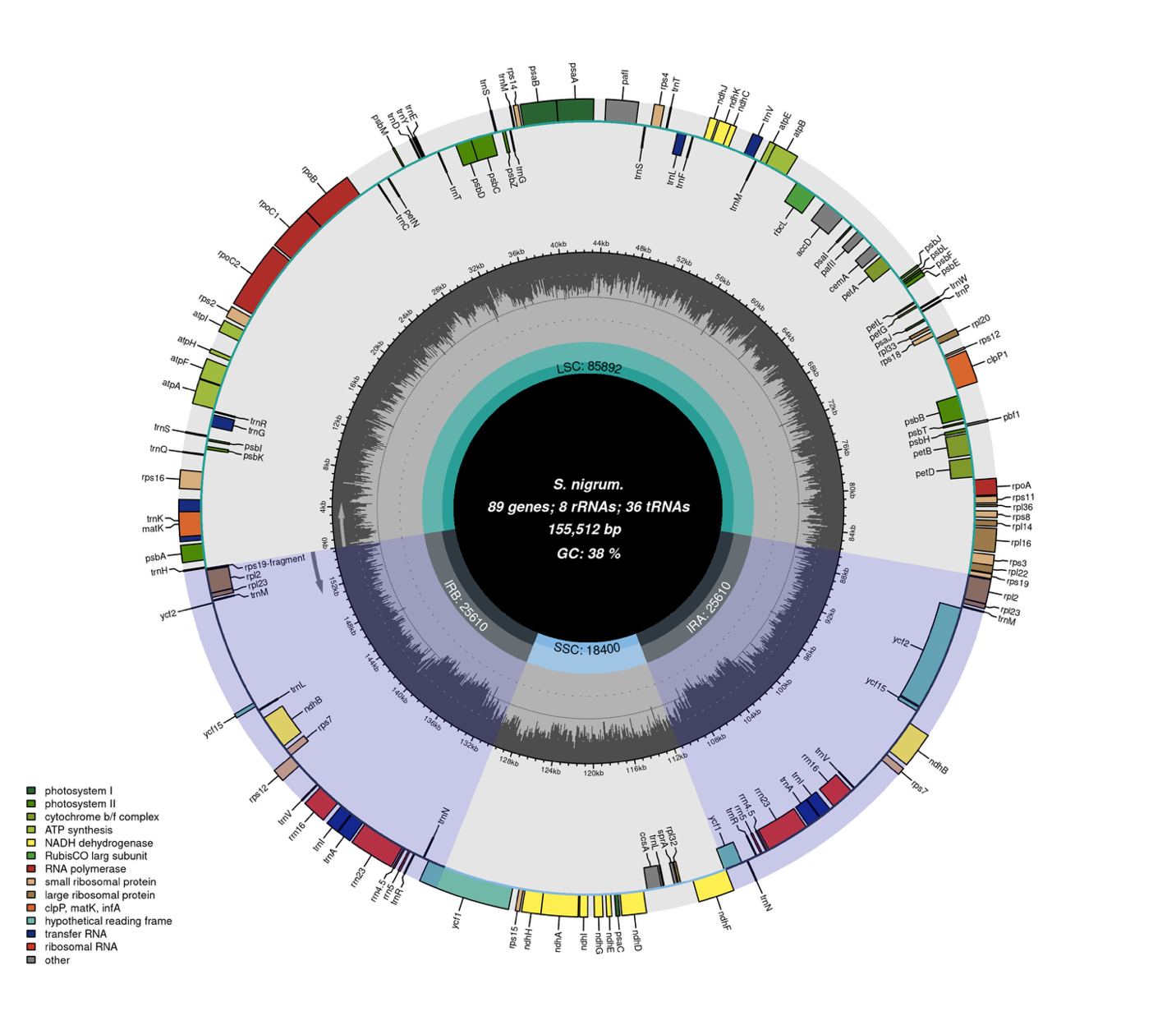

**Supplementary Figure 1: Gene map of the representative circular chloroplast genome of *S. nigrum* (PV802130).** Annotations were created with GeSeq and manually adjusted in Geneious. The plastome was visualized using Chloroplot. The innermost circle describes species name, gene counts, plastome length, and GC content. The inner grey circle represents the variation in GC content across the genome. The two IR regions are highlighted in blue. The outer circle shows the gene names coloured by their function. Inner genes are transcribed in a clockwise direction, while outer genes are transcribed in a counterclockwise direction.

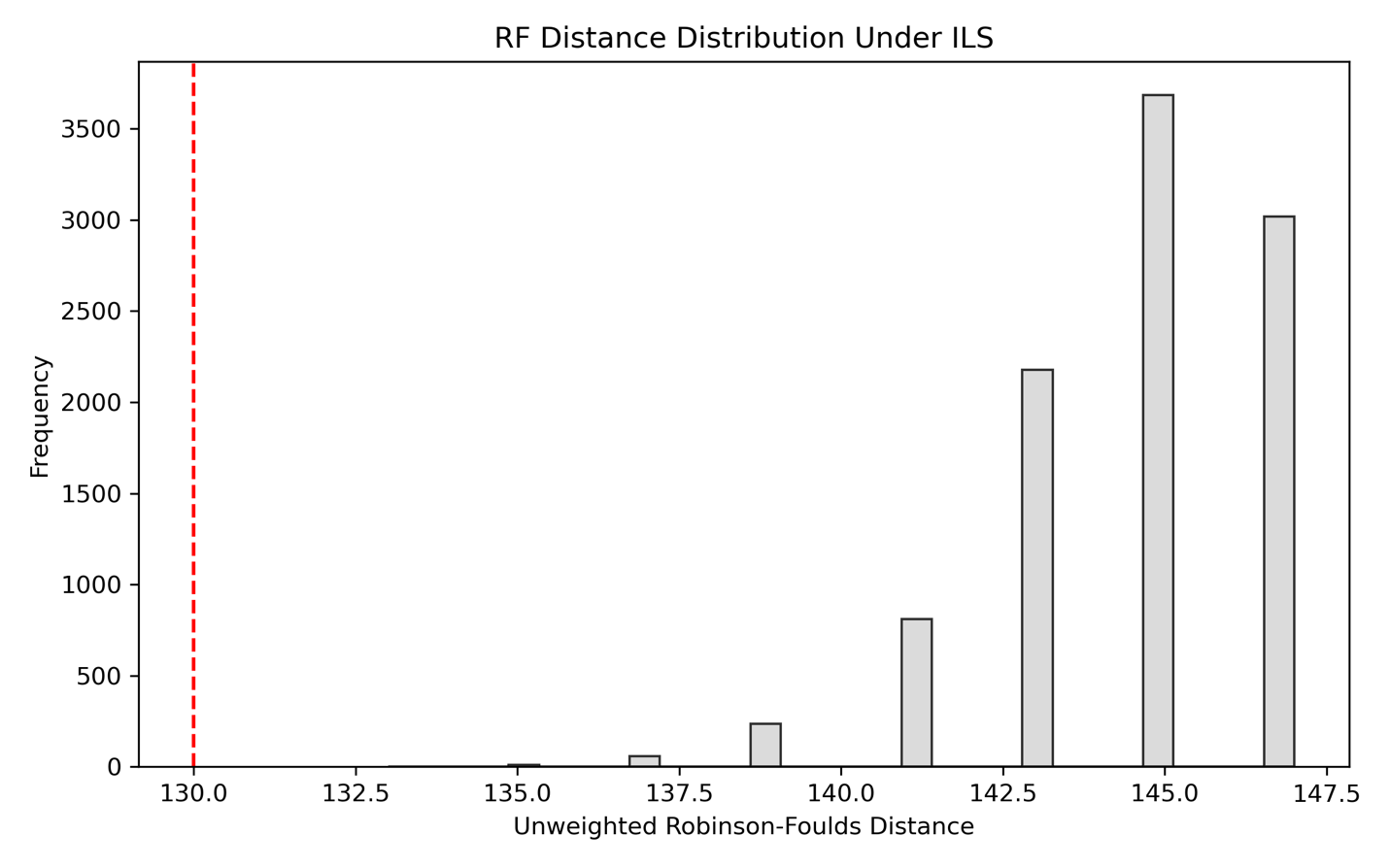

**Supplementary Figure 2: Distribution of RF distances from ILS simulations (4× scaling) for plastome gene trees versus the nuclear tree**. The red dashed line marks the observed RF=130 between our empirical plastome and nuclear phylogenies. The grey bars represent the distribution of RF distances when simulated.
